# Distinct developmental changes in linear and nonlinear neural interactions across infancy and adulthood

**DOI:** 10.1101/2025.08.18.670844

**Authors:** Lorena Santamaria, Stanimira Georgieva, Valdas Noreika, Andres Canales-Johnson, Victoria Leong

## Abstract

The development of functional brain connectivity during early life depends on social experience and is best understood in the context of interactions with a caregiver. It is still an open question to what extent distinct modes of functional connectivity dominate in different stages of human brain development, and whether cognitive tasks can differentially modulate them. Using electroencephalography (EEG), we investigated the development of linear and nonlinear functional brain connectivity in infants and adults while they socially interacted. Using simultaneous EEG recordings in parent-infant dyads performing two different experiments (N = 160; 80 adults and 80 infants), we computed functional connectivity capturing distinct dynamics: a linear measure capturing phase synchronization (weighted phase lag index; WPLI), and a nonlinear measure capturing information sharing (weighted symbolic mutual information; WSMI). In both tasks, adults showed higher WSMI than infants, whereas infants showed higher WPLI than adults. Moreover, infant age predicted only task-related connectivity values computed with the nonlinear measure and not with its linear counterpart, suggesting that the information-theoretic measure was more sensitive to developmental changes in task-relevant neural processing. These findings suggest that over development, a shift in dominance from linear to nonlinear modes of brain communication may be essential for supporting emerging higher cognitive abilities, such as precursors to executive function (here, attention shifting) and social decision-making. Further, this work highlights the importance of using nonlinear measures in addition to traditional linear ones, which collectively permit a more robust capture of maturational changes.

## INTRODUCTION

The development of functional brain connectivity during early life depends on social experience and is best understood in the context of scaffolding interactions with a caregiver (Endevelt-Shapira and Feldman, 2023; Ilyka et al., 2021; Li et al., 2023; Perone and Gartstein, 2019; Reindl et al., 2018). Brain networks follow a maturation trajectory in the first 2 years of life, from primary sensory and motor systems present prenatally (Moore et al., 2023) to complex language and reasoning centres, accompanied by changes in local and long-range functional connectivity (Bosch-Bayard et al., 2022; Cao et al., 2016; Gao et al., 2011; Gilmore et al., 2018; Khundrakpam et al., 2016). Early social interaction actively shapes the functional developmental process with a long-lasting impact (Ilyka et al., 2021; Perone and Gartstein, 2019; Tan et al., 2023). These rapid developmental changes in brain maturation contribute to individual differences in neural dynamics in response to varying task demands. Developmentally significant changes in functional connectivity have been shown in relation to sensori-motor processing (Allievi et al., 2015), fine and gross motor skills (Bruchhage et al., 2020; Marrus et al., 2017), visual perception (Caffarra et al., 2023; Wilcox et al., 2012), and visual temporal processing (Freschl et al., 2022), auditory scene perception (Nakano et al., 2009), social skills (Eggebrecht et al., 2017; Grossmann and Johnson, 2007), and language ability (Dehaene-Lambertz et al., 2002; Perani et al., 2011). Using fMRI recordings during sleep, Bruchhage et al. (2020) investigated the correlation between functional connectivity and Mullen Scales of Early Learning in 196 typically developing children between 3 months and 6 years of age. They found positive correlations between specific networks (anatomical areas) and visual reception, motor skills (fine and gross) and language after correcting by age (Bruchhage et al., 2020).

Changes in functional connectivity are believed to represent shifts in the balance of information segregation and integration (Liao et al., 2017). During the first 2 years of life, the maturing brain changes from a relatively randomized configuration to a well-organized one with progressively higher robustness, global clustering, and efficiency (Cao et al., 2016; Damaraju et al., 2014; Gao et al., 2011). While modularity may decrease, local and global efficiency increases with age (Boersma et al., 2011; Richmond et al., 2016). The maturation process involves a combination of synaptic growth and myelination, among others, which occur at micro- and macroscopic levels. Most of these processes follow an asynchronous course. For instance, myelination of the white matter pathways at the macroscopic level progresses along a posterior-to-anterior topographical sequence (Sotardi et al., 2021). White matter asymmetry has also been reported with earlier maturation in the left hemisphere for the frontal region (Sotardi et al., 2021). Recent advances in structural connectivity also revealed that neural tracts organisation follows a nonlinear modification with age (Dubois et al., 2014; Lebel and Beaulieu, 2011). Similarly, nonlinear development of EEG connectivity has been reported (Lochy et al., 2019), with earlier connectivity peaking for local occipital and parietal, and later in frontal networks (Bosch-Bayard et al., 2022), while long-range functional connectivity increases with age (Falivene et al., 2024), and in relation to cognitive ability (Falivene et al., 2024; Ramos-Loyo et al., 2022).

Traditional measures of functional connectivity (e.g., phase coherence or Granger causality) primarily quantify interactions within the same frequency band and are mainly sensitive to linear signal transmission (Pesaran et al., 2018; Schneider et al., 2021; Vinck et al., 2025, 2023). Such linear dependencies may simply reflect anatomical feedforward projections from one region to another (Dowdall et al., 2023; Dowdall and Vinck, 2023; Schneider et al., 2021), rather than functional integration (Vinck et al., 2023). In contrast, growing evidence suggests that neural communication often relies on nonlinear, cross-frequency, and arrhythmic dynamics that these conventional measures cannot capture (Imperatori et al., 2019; Vinck et al., 2025, 2023). In recent years, this has motivated a shift towards information-theoretic approaches to study brain interactions (Blume et al., 2024; Canales-Johnson et al., 2023; Chidichimo et al., 2025; Ince et al., 2017; Roberts et al., 2025) that capture nonlinear transformation across regions, independent of oscillatory synchrony or spectral overlap (Vinck et al., 2025, 2023). Nonlinear functional connectivity measures, such as weighted symbolic mutual information (WSMI) (King et al., 2013), quantify information sharing between neural signals, indicating nonlinear coupling of task-related variables, such as those observed during social interactions. WSMI has revealed neural coordination patterns that linear analyses often miss (Canales-Johnson et al., 2020b; Imperatori et al., 2019; King et al., 2013; Olivares et al., 2025; Potash et al., 2025; Sitt et al., 2014), and has proven particularly valuable in capturing transient, task-specific interactions across distributed brain networks (Canales-Johnson et al., 2020a,b).

Although brain functional connectivity can be detected from an early age, its contribution to cognitive processes during development is not well understood. For example, several electroencephalography (EEG) studies have emphasized the relevance of alpha and theta (3-9 Hz) neural synchronization during infant-adult social interactions (Endevelt-Shapira and Feldman, 2023; Leong et al., 2017; Perone and Gartstein, 2019; Santamaria et al., 2020; Wass et al., 2018). However, infant and adult EEG exhibits a diverse spectrum of neural dynamics, i.e., not only oscillatory activity in specific frequency ranges, but also aperiodic activity devoid of a particular temporal beat (non-oscillatory activity). For example, significant developmental changes have recently been reported in slope and offset of the aperiodic component of EEG (Cellier et al., 2021; Donoghue et al., 2020; Wilkinson et al., 2024). Longitudinal studies extending from childhood to adulthood have observed decreases in aperiodic slope with age (Hill et al., 2022; McSweeney et al., 2023; Merkin et al., 2023; Voytek et al., 2015). The functional significance of these developmental changes in the aperiodic component is unclear in relation to functional connectivity and cognitive processes. However, since infant brains during development tend to exhibit a high degree of heterogeneity in frequencies across areas (Cellier et al., 2021), it is implausible, given the almost all-to-all anatomical connectivity in the cortex, that integration of task-related activity can rely solely on synchronization within the same frequency band. Possibly, varying task demands would interact with age-related changes in cortical maturation to produce differential sensitivity of cognitive processes in nonlinear functional connectivity.

To address these issues, here we used simultaneous EEG recordings in infant-adult dyads performing two different experimental tasks. We first compare linear (i.e., weighted phase lag index, or WPLI) and nonlinear (i.e., WSMI) functional connectivity in developing (infant) and mature (adult) neural systems (Figure 1). Second, we investigate dynamic and task-relevant modulation of each form of functional connectivity in a socially interactive context.

**Figure 1:**
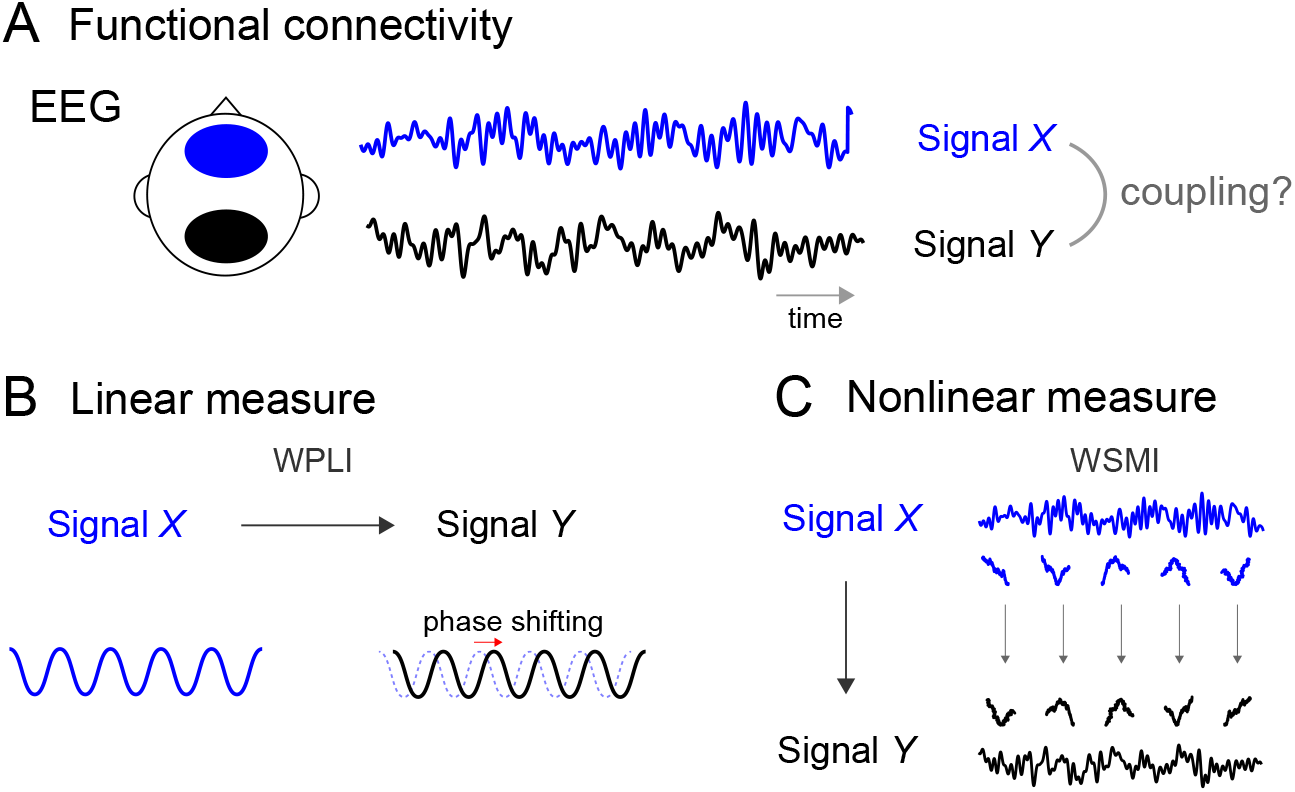
Experimental design, linear and nonlinear functional connectivity. **(A)** Mapping neural connectivity between two EEG signals. In this diagram, the mapping (functional connectivity) across two signals is depicted between signal X (posterior electrodes) and signal Y (anterior electrodes). **(B)** Mapping functional connectivity between signal X and Y. Linear transformations between two correlated signals can only occur at the same frequency; that is, the original signal (signal X) can only result in a scaled or phase-shifted version of the signal (signal Y) at the same frequency (e.g., a 10 Hz signal X can result in a phase-shifted or amplitude-increase signal Y at the same 10 Hz). The WPLI measure captures this type of linear mapping (weighted phase lagged index). **(C)** While linear measures are necessarily blind to relationships occurring across different frequencies, nonlinear transformations cause systematic relationships across different frequencies between signal X and Y. Nonlinear functional connectivity measures quantify arbitrary mappings between temporal patterns in one signal (signal X) and another signal (signal Y). This type of nonlinear mapping is captured by the WSMI measure (weighted symbolic mutual information).

## RESULTS

### Experiment 1: Affective Decision Making Task

To understand the dynamic processing of affective information in infant and adult brains while engaged in a social interaction, we first investigated linear and nonlinear functional brain connectivity in the dyads performing the Affective Decision Making (ADM) Task. Functional connectivity was computed on temporal windows corresponding to the Presentation period (mother expressed her like and dislike for each object while the infant observed; see Methods); the Response period (infant selected one of the two objects); and the Interval (infant interacted freely with the objects).

Thus, we performed a repeated measures ANOVA (RANOVA) with three factors: Stage (Infant vs. Adult), Dynamics (WPLI vs. WSMI), and Period (Presentation vs. Response vs. Interval). We observed a significant main effect of Dynamics (F_(1,46)_= 679.35; p < 0.001; *η*^2^*p* = 0.93) and Period (F_(1,46)_ = 21.29; p < 0.001; *η*^2^*p* = 0.31), but no effect for Stage (F_(1,46)_ = 0.03; p < 0.98). (Figure 2A).

**Figure 2:**
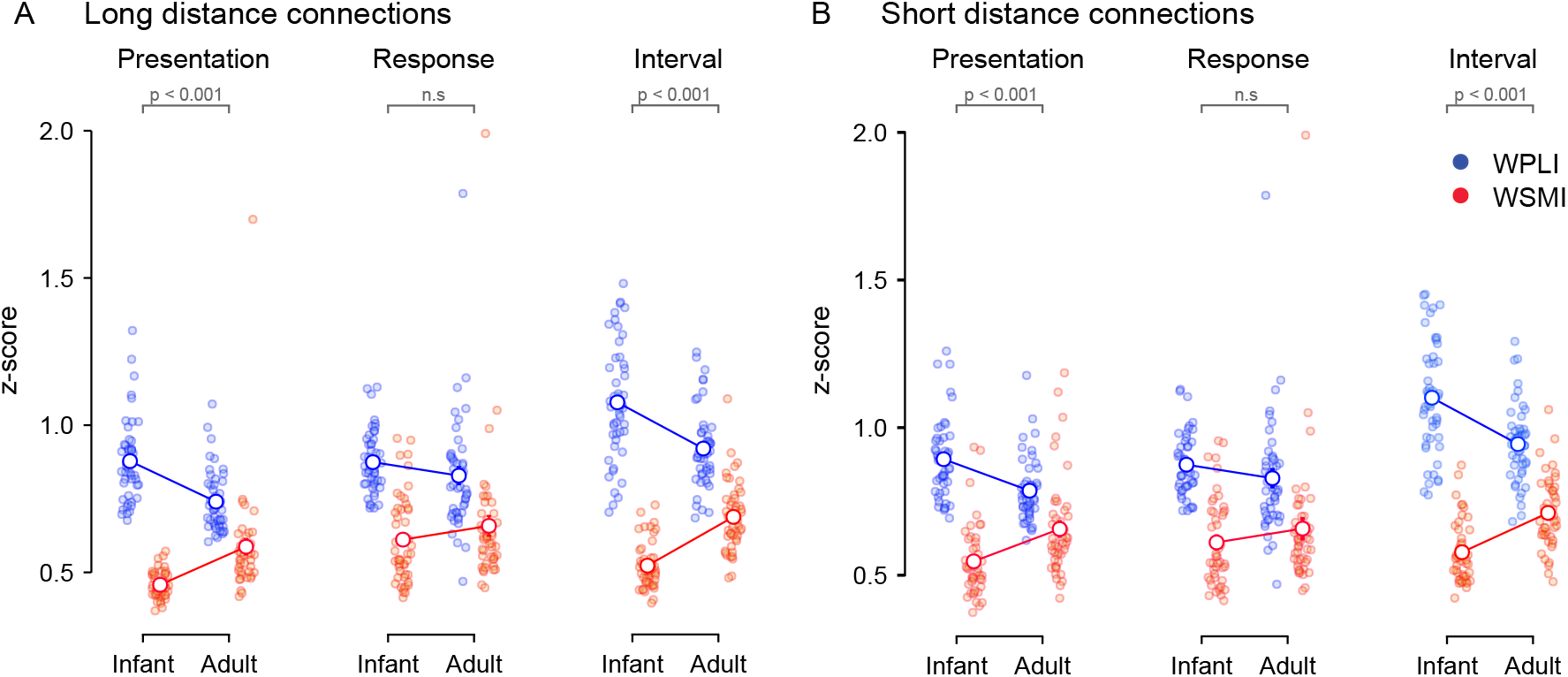
Linear and nonlinear functional connectivity between Adults and Infants in the ADM task. **(A)** Long-distance connectivity maps (Higher 40% of the connections (in cm) between electrodes) for the Presentation, Response, and Interval period during ADM Task. Individual dots represent the mean WSMI (nonlinear functional connectivity; in red) and WPLI (linear functional connectivity; in blue) values across the corresponding EEG electrode pairs for each participant (i.e., 47 Infants and 47 Adults). The white dot represents the mean across participants. Error bars represent the standard error of the mean (S.E.M). To make WSMI and WPLI comparable in frequency content, connectivity was computed using a common frequency range captured by both metrics (< 16 Hz). To make the results comparable for further statistical analyses, results were normalized per participant using z-scores. **(B)** Short-distance connectivity maps (Lower 40% of the connections (in cm) between electrodes) for the Presentation, Response, and Interval period during ADM Task. All details are the same as in (A).

In the case of long-distance connections (i.e., Higher 40% of the connections measured in centimeters between electrodes), we observed a significant triple interaction between Stage, Dynamics, and Period (F_(1,46)_ = 16.74; p < 0.001; *η*^2^*p* = 0.26). Interestingly, post-hoc comparisons showed an increase in WSMI in Adults compared to Infants (t_(1,46)_ = 4.70; p < 0.001) and a decrease in WPLI in Adults compared to Infants in the Presentation period (t_(1,46)_ = −5.42; p < 0.001). Similarly, during the Interval, adults showed higher WSMI compared to infants (t_(1,46)_ = 10.98; p < 0.001), but lower WPLI compared to Infants (t_(1,46)_ = 8.34; p < 0.001). However, no differences between WPLI and WSMI during the Response period (WSMI Adult vs. Infant: t_(1,46)_ = 1.21; p = 0.690; WPLI Adult vs. Infant: t_(1,46)_ = −1.64; p = 0.827).

For short-distance connections (i.e., lower 40% of the connections measured in centimeters between electrodes), we observed a significant main effect of Dynamics (F_(1,46)_= 546.19; p < 0.001; *η*^2^*p* = 0.92), and Period (F_(1,46)_= 17.86; p < 0.001; *η*^2^*p* = 0.28), but no effect for Stage (F_(1,46)_ = 0.05; p = 0.808; *η*^2^*p* = 0.001). We observed a significant triple interaction between Stage, Dynamics, and Period (F_(1,46)_ = 13.17; p = 0.001; *η*^2^*p* = 0.22). Again, in the Presentation period, post-hoc comparisons showed an increase in WSMI in Adults compared to Infants (t_(1,46)_ = 3.76; p = 0.011) and a decrease in WPLI in Adults compared to Infants in the Presentation period (t_(1,46)_ = −4.28; p = 0.002). The Interval showed an increase in WSMI (t_(1,46)_ = 8.11; p < 0.001) but decreased WPLI in Adults compared to Infants (t_(1,46)_ = −8.22; p < 0.001). Finally, no differences were found between WSMI and WPLI during the Response period (WSMI Adult vs. Infant: t_(1,46)_ = 1.21; p = 1.000; WPLI Adult vs. Infant: t_(1,46)_ = −1.46; p = 1.000). (Figure 2B).

Together, these results indicated that the neural connectivity underpinning engagement in affect communication was more readily described by the nonlinear measure in the adult brain and the linear one in the infant brain. This dissociation, however, was dynamically modulated by the task period and was not present during the infant’s response window.

### Experiment 2: Sequential Touching Task

Next, we investigated linear and nonlinear functional connectivity in infant-adult dyads engaged in an attentional set-shifting sequential touching task (STT) (Tan and Leong, 2023). In STT, functional connectivity measures were computed on temporal windows corresponding to periods of infant’s free object play before (Pre) and after (Post) mother demonstrated the less salient property of the objects (Demo, see Methods). Thus, we performed a repeated measures ANOVA (RANOVA) with three factors: Stage (Infant vs. Adult), Dynamics (WPLI vs. WSMI), and Period (Pre vs. Demo vs. Post).

In the case of long-distance connections, we observed a significant main effect of Stage (F_(1,46)_= 19.78; p < 0.001; *η*^2^*p* = 0.38), Dynamics (F_(1,46)_= 1725.47; p < 0.001; *η*^2^*p* = 0.98), and Period (F_(1,46)_ = 4.02; p = 0.023; *η*^2^*p* = 0.11). Crucially, we observed a significant triple interaction between Stage, Dynamics, and Period (F_(1,46)_ = 26.38; p < 0.001; *η*^2^*p* = 0.45). In the Pre period, post-hoc comparisons showed an increase in WSMI in Adults compared to Infants (t_(1,46)_ = 5.11; p < 0.001) and a decrease in WPLI in Adults compared to Infants in the Pre period (t_(1,46)_ = 12.28; p < 0.001). Similarly, in the Post period, we observed increased WSMI in adults compared to Infants (t_(1,46)_ = 4.48; p = 0.001), and a decrease in WPLI between Adults and infants (t_(1,46)_ = 10.84; p < 0.001)(Figure 3A). However, no differences were found between WPLI and WSMI during the Demo period (WSMI Adult vs. Infant: t_(1,46)_ = 1.44; p = 1.00; WPLI Adult vs. Infant: t_(1,46)_ = −1.43; p = 1.00).

**Figure 3:**
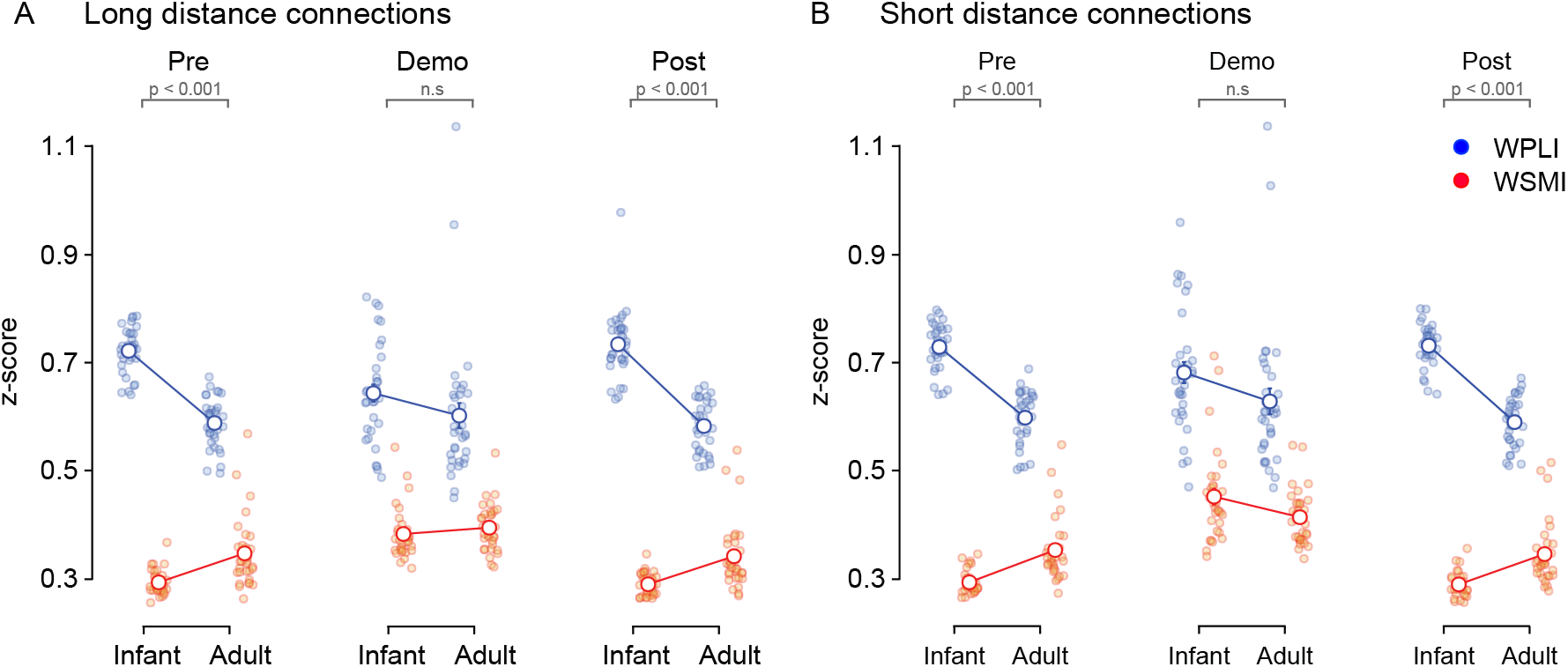
Linear and nonlinear functional connectivity between Adults and Infants in the STT. **(A)** Long-distance connectivity maps (Higher 40% of the connections (in cm) between electrodes) for the Pre, Demo, and Post period during STT. Individual dots represent the mean WSMI (nonlinear functional connectivity; in red) and WPLI (linear functional connectivity; in blue) values across the corresponding EEG electrode pairs for each participant (i.e., 33 Infants and 33 Adults). The white dot represents the mean across participants. Error bars represent the standard error of the mean (S.E.M). To make WSMI and WPLI comparable in frequency content, connectivity was computed using a common frequency range captured by both metrics (< 16 Hz). To make the results comparable for further statistical analyses, results were normalized per participant using z-scores. **(B)** Short-distance connectivity maps (Lower 40% of the connections (in cm) between electrodes) for the Response and Task period during STT. All details are the same as in (A).

For short-distance connections, we observed a significant main effect of Stage (F_(1,46)_= 21.71; p < 0.001; *η*^2^*p* = 0.40), Dynamics (F_(1,46)_= 1887.17; p < 0.001; *η*^2^*p* = 0.98), and Period (F_(1,46)_ = 22.49; p < 0.001; *η*^2^*p* = 0.41). Again, we observed a significant triple interaction between Stage, Dynamics, and Period (F_(1,46)_ = 37.74; p < 0.001; *η*^2^*p* = 0.54). In the Pre period, post-hoc comparisons showed an increase in WSMI in Adults compared to Infants (t_(1,46)_ = 5.87; p < 0.001) and a decrease in WPLI in Adults compared to Infants in the Pre period (t_(1,46)_ = 11.86; p < 0.001). Similarly, in the Post period, we observed increased WSMI in adults compared to Infants (t_(1,46)_ = 5.14; p = 0.001), and a decrease in WPLI between Adults and infants (t_(1,46)_ = 12.57; p < 0.001). However, there were no differences between WPLI and WSMI during the Demo period (WSMI Adult vs. Infant: t_(1,46)_ = 2.70; p = 0.109; WPLI Adult vs. Infant: t_(1,46)_ = −1.69; p = 0.598). (Figure 3B).

Similar to the ADM task, when dyads engaged in the attentional flexibility task, adult neural functional connectivity showed an increase in the nonlinear measure, in contrast to infant connectivity, which had an increase in the linear measure. Again, this dissociation changed dynamically depending on the task period.

### Infant’s age predicts nonlinear functional connectivity

We further investigated the statistical dependencies between the infant’s age and linear and nonlinear functional connectivity. Separate multivariate multiple regressions were performed on the ADM and STT. In ADM, we used WPLI and WSMI as regressors in the Presentation and Interval. In STT, the predicted variables were WSMI and WPLI values in the Pre and Post periods. For each task, the periods mentioned above were selected due to the significant modulation of WSMI and WPLI reported between infants and adults in the previous section. For both regressions, the age of the infants (in weeks) was used as the regressor variable. In the ADM task model (Wald χ^2^(2) = 8.24, p = 0.0163), infant’s age predicted WSMI in the Interval (t_(45)_ = 2.124, p = 0.0392) but not in the Presentation period (t_(45)_ = 1.837, p = 0.0728). In the case of WPLI, the model was not significant for either period (Wald X^2^(2) = 0.45, p = 0.7984). In the STT model (Wald χ^2^(2) = 7.49, p = 0.0236), infant’s age predicted WSMI in Post (t_(31)_ = 2.583, p = 0.0147) but not in Pre (t_(31)_ = 1.108, p = 0.2763). In the case of WPLI, the model was not significant (Wald χ^2^(2) = 4.43, p = 0.1090) for either period. These relationships are depicted as simple Pearson correlations in Figure 4.

**Figure 4:**
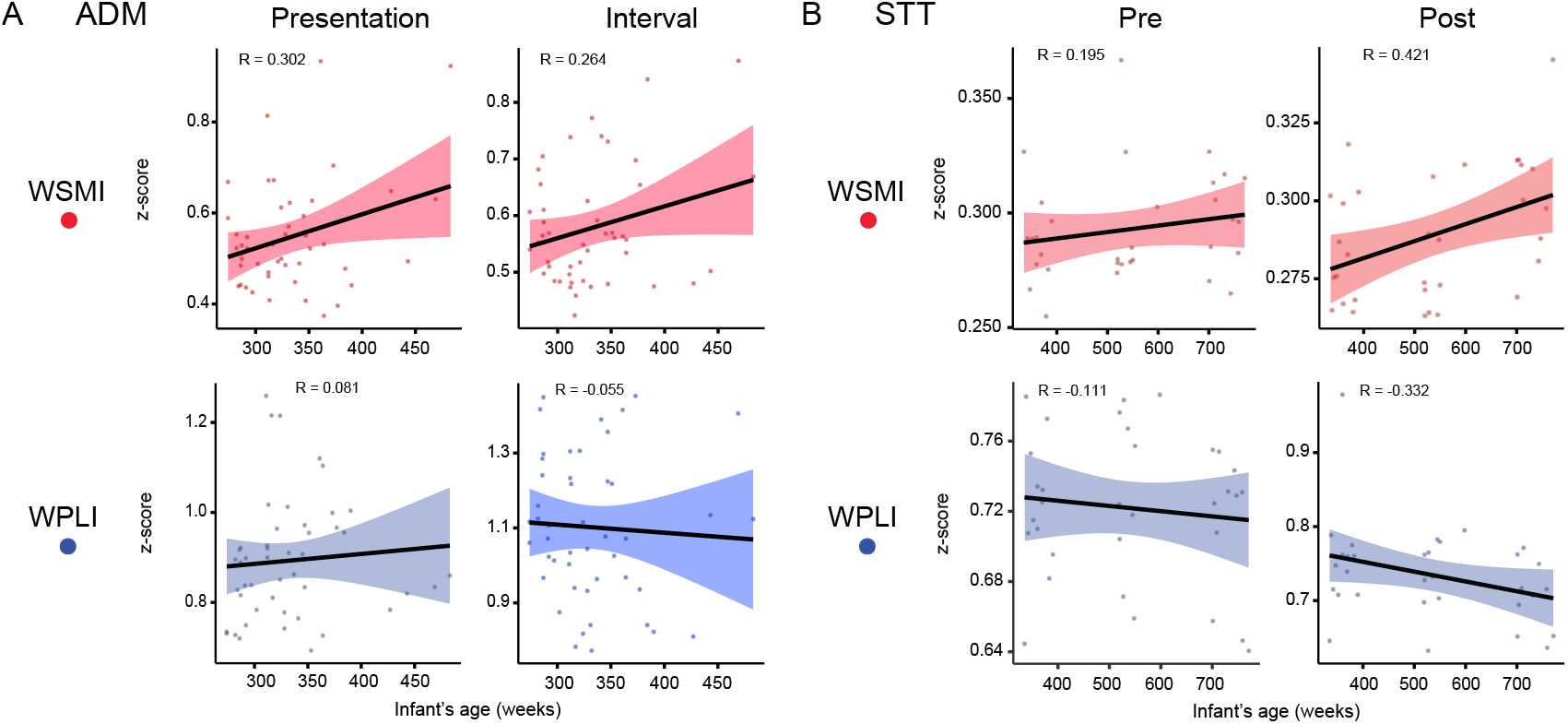
Correlations between infant’s age and WSMI and WPLI. **(A)** Pearson’s correlations in the Presentation and Interval conditions for the ADM task. Z-scored WSMI (red) and WPLI (blue) functional connectivity values are depicted against the infant’s age (in weeks). **(B)** Pearson’s correlations in the Pre and Post conditions for the STT. Z-scored WSMI (red) and WPLI (blue) functional connectivity values are depicted against the infant’s age (in weeks).

### Brain-behavior relationships

We further investigated the statistical dependencies between WPLI and WSMI, and the amount of attentional shifting infants showed during STT, as measured by shiftQ (see Methods for details). Two multiple regressions were used to predict the ShiftQ index from the long-distance and short-distance connections in Infants. The predictive variables in each model were WSMI and WPLI in the Pre, Demo, and Post periods, as well as Age. In the long-distance connections model (R^2^ = 0.40; F_(7,25)_ = 2.47; p = 0.045), only WPLI in the Post condition positively predicted ShiftQ (β = 1.16; p = 0.025). In the case of the short-distance connections model (R^2^ = 0.44; F_(7,25)_) = 2.85; p = 0.025), only WSMI in the Pre condition positively predicted ShiftQ (β = 1.90; p = 0.037). The results indicate that in infants, the amount of attentional shifting increases when long-distance linear connectivity is lower after demonstration, but local nonlinear connectivity is lower before demonstration.

## DISCUSSION

Here, we investigated the extent to which linear and non-linear functional brain connectivity dominate during different stages of human development and whether cognitive demands differentially modulate the recruitment of these processes. Using simultaneous EEG recordings in parent-infant dyads performing two separate experiments (N = 160; 80 adults and 80 infants), we computed functional connectivity metrics measuring linear (WPLI) and nonlinear (WSMI) neural interactions. We compared the linear and nonlinear dynamics in developing (infant) and mature (adult) neural systems and assessed the dynamic modulation of each form of connectivity in a socially interactive context in the two tasks.

In both experiments, we observed that a dissociation between WPLI and WSMI characterized the infant-mother dyad during their interactions in the task. While WSMI increased in adults compared to infants, WPLI showed the opposite pattern. These results support a greater reliance on nonlinear dynamics in the mature brain, and conversely, a greater reliance on linear dynamics in the developing brain. Interestingly, we observed a dynamic modulation in the two patterns of connectivity, whereby both tasks contained a period that did not show a linear-nonlinear dissociation between adults and infants. These task periods were characterised by specific cognitive demands on the infant: in ADM, they performed affective decision making in the context of maternal social referencing; in STT, they controlled and shifted their attention between salient and non-salient object feature dimensions.

### Functional roles of nonlinear and linear neural interactions

Although the mainstream view is that cognition modulates functional connectivity between oscillations at the same frequency (Fries, 2015), other perspectives emphasize the non-linear nature of neural communication, which may require non-oscillatory rather than oscillatory dynamics (Singer, 2021; Vinck et al., 2024, 2023). In this alternative view, cognitive processing modulates nonlinear connectivity as a consequence of interactions across frequency bands rather than interactions within the same frequency band, as is the case with linear functional connectivity. This idea is based on evidence demonstrating that nonlinear interactions are functionally relevant for complex computations such as pattern extraction in neural networks, leading to the maturation of executive functions such as attention (Vinck et al., 2023), and for the long-range communication of perceptual and predictive information (Canales-Johnson et al., 2020a,b; Gelens et al., 2024; Potash et al., 2025; Vinck et al., 2023).

During the development of the cortex, it is plausible that functional connectivity across cortical regions with different maturation trajectories may differ in frequency ranges (Bosch-Bayard et al., 2022) and thus may not be readily captured by linear but by nonlinear functional interactions.

### Linear and nonlinear functional connectivity changes between infants and adults

The functional balance of linear and nonlinear connectivity processes may change from infancy to adulthood due to the dynamic nature of brain maturation. More basic sensory and motor processing may rely more readily on linear connectivity. In contrast, higher cognitive functions, which entail increased information integration, may be better captured by nonlinear connectivity measures (Canales-Johnson et al., 2020a, 2023, 2021b; Potash et al., 2025; Vinck et al., 2023). In infants, anatomical brain connectivity is dominated by short-range and localized patterns, with relatively immature myelination, synaptic pruning, and dendritic arborization. This developmental state can limit the complexity and efficiency of long-range communication, resulting in functional connectivity patterns that are predominantly simpler and more linear in nature (Gilmore et al., 2018; Menon, 2013; Oldham and Fornito, 2019; Oldham et al., 2022).

As the brain matures, synaptic plasticity, network specialization, and integration enhance long-range connectivity, allowing more intricate and flexible communication. These changes foster the emergence of nonlinear interactions, which might better capture the complex dynamics of cross-frequency coupling of mature brain function. The development of higher cognitive functions and adaptive behavior in adults is based on a shift towards efficient global integration, which inherently involves nonlinear mechanisms (Vinck et al., 2023). For example, non-linear interactions have been shown to capture better complex pattern extraction in neural networks relevant for the development of high-order executive functions such as attention (Vinck et al., 2023), cognitive control (Canales-Johnson et al., 2020a), and perceptual integration and segregation during ambiguous perception (Canales-Johnson et al., 2023, 2020b). While the maturation trajectory of nonlinear dynamics about brain function from infancy to adulthood is not yet well known, functional connectivity measures in infancy show increased complexity and efficiency (Boersma et al., 2011; Cao et al., 2016; Damaraju et al., 2014) and a significant reorganisation in adolescence (Gabard-Durnam et al., 2014; Uhlhaas et al., 2009). These changes have been linked to developmental sophistication in both cognitive and affective processes (Kar et al., 2013; Uddin et al., 2011). These developmental differences reflect the brain’s structural and functional evolution, which adapts connectivity patterns to meet the growing cognitive and behavioral demands of adulthood.

In conclusion, these findings suggest that over development, a shift in dominance from linear to nonlinear forms of brain communication may be essential for supporting emerging higher cognitive abilities, such as precursors to executive function (here, attention shifting) and social decision-making. Further, this work highlights the importance of using nonlinear measures in addition to traditional linear ones, which collectively permit a more robust capture of maturational changes.

## METHODS

### Affective Decision-Making Task (ADM)

#### Participants

Forty-nine mother-infant dyads participated in the experiment, of which 22 were male and 27 female. Two participants were rejected for not producing sufficient usable trials (fewer than 2), so 47 dyads remained in the analysis. The average age of the infants was 335.95 days (±7.29 days, standard error of the mean). All mothers reported no neurological problems and normal hearing and vision for themselves and their infants. A written consent form was collected from all subjects. The protocol was approved by the Cambridge Psychology Research Ethics Committee (PRE.2016.0.29).

#### Materials

Four pairs of novel objects were used. The objects within each pair were matched in size and texture but had different shape and colour. The order of object presentation (positive or negative first) was counterbalanced across trials, and the side of the object presentation (e.g., positive on the left or right) was also randomised and counterbalanced across trials within each participant. That is, the last object presented was positive 50% of the time and negative 50% of the time. Similarly, positive objects were presented on the left 50% of the time, and on the right 50% of the time (and vice versa for negative objects). Further, the order and side of presentation were changed for each successive trial (while retaining the valence marking within object pairs). Finally, the order of presentation of the four pairs of objects was counterbalanced across participants.

#### Task

We used an affective decision-making task involving positive and negative maternal emotional demonstrations towards novel toy objects (Santamaria et al., 2020). Infants were seated in a high chair, and a table was placed immediately in front of them. Parents were seated on the opposite side of the table, directly facing the infant. The width of the table was 65cm.

A trial began when the mother attracted her infant’s attention by saying “Look”, or by holding one of the objects up. Each experimental trial comprised a maternal presentation period involving one pair of novel objects, a response period, and an interval. During the Presentation period, mothers were instructed to show positive affect toward one object and negative affect toward the other object. Mothers were instructed to limit their speech to simple formulaic verbal statements per object (which they repeated for each object), and to model positive or negative emotions in a prescribed manner (e.g., smiling versus frowning). The start (onset) of the Presentation period of each trial was determined as the point when the mother began speaking about either the positive or negative object, and was completed when maternal utterances ended. (Santamaria et al., 2020). The Response period started at the end of the maternal presentation and ended with the infant touching one or both of the objects. Next, infants were allowed to interact briefly with the objects before retrieving them (the Interval). The start and end points of each task period were determined by manual video coding.

Each of the four pairs of objects was presented four times to each infant, yielding four sets of four trials and a maximum of sixteen possible trials in total. The task was discontinued if infants showed prolonged fussiness or inattention. An experimenter was present throughout the session, but positioned out of the line of sight of both participants, to ensure the participants were interacting as instructed. The experimenter provided new pairs of objects as required and informed the parent regarding the side and order (pos/neg) of presentation for each pair of items before the start of each trial, but explicitly avoided making prolonged social contact with either participant.

### Sequential Touching Task (STT)

#### Participants

Fifty-two mother-infants participated in the study. From this cohort, 7 participants were discarded due to technical difficulties (no EEG files or EEG markers) and 8 further participants were discarded due to low EEG quality, leaving a total of 39 usable dyads (15 male infants). 6 participants were rejected for not producing sufficient usable trials (fewer than 2). At the time of the experiment, the age of the infants was 426 days (±22.68) on average. All mothers and infants were healthy volunteers with no neurological problems. The study was approved by the Nanyang Technological University Institutional Review Board (IRB-2021-808).

#### Materials

Eight objects were selected for this task with two distinctive physical properties: shape (4 balls vs 4 cubes, high salience) and material (4 hard vs 4 soft, low salience). All objects had the same size but differed in colour to retain the interest of the infant in the toys (same materials were used as depicted in Tan and Leong (2023)).

#### Task

Mothers and infants performed a social interactive attention set-shifting task suitable for infants aged 12-24 months (Tan and Leong, 2023). The task comprised three parts lasting a total of ~10 min: two periods of free play with the objects (~4 min each) before and after a demonstration period (~2 min). During the free play (Pre and Post) periods, the infant was allowed to interact freely with the full set of toys, with minimal interaction from the mother. During the Demonstration (Demo) period, the mother was seated in front of the infant and compressed each toy one at a time whilst her infant watched, to flag the less salient property of each object, i.e., how soft or hard the material was. The toy demonstration order was randomised across participants. The task tested how infants shifted their attention towards the more or less salient properties of the objects following the demonstration.

### EEG acquisition and pre-processing

For the STT, EEG was recorded using a 32-channel LiveAmp amplifier (Brain Vision©) acquiring 30 channels of EEG and 2 channels of ECG. The ground was placed on the forehead while Cz was used as a reference. For the ADM task, the EEG was acquired using a 32-channel BIOPAC Mobita mobile amplifier for both infant and adults. The ground electrode was affixed to the back of the neck, and reference was done offline. In both experiments, an Easycap electrode system was used for both infants and adults, with electrode placement following the international 10/20 system. Electrode impedance was kept below 10KΩ for adults and 20KΩ for infants, and the sampling rate was 500 Hz.

For both tasks, pre-processing was performed using the EEGlab toolbox (Delorme and Makeig, 2004) and custom Matlab© functions (as in many of our previous studies (e.g., Lanfranco et al. (2021)). Data were low-pass and high-pass filtered (40Hz and 0.5Hz, respectively). Bad channels were visually detected and set as empty (NaNs), participants with 7 or more noise channels were not included for further analysis. Furthermore, eye and muscle-related artefacts were removed using independent component analysis (ICA) (Delorme and Makeig, 2004). An IC component was removed from the adult EEG only if it could be attributed to a clear, specific noise source: eye blinks, saccades, or muscle. For the infants, a similar approach was followed, but fewer eye-related ICs were detected. The data were re-referenced to a common average reference, and Cz was added back, giving a total of 31 (STT) and 32 (ADM) channels. Furthermore, highly noisy segments that were still present after ICA pruning were also replaced with NaNs in both infants’ and adults’ EEG recordings to keep the time structure of the data.

### Attentional Set-shifting metric: ShiftQ

ShiftQ is a measure of the shift in an infant’s object categorisation strategy from pre-play to post-play, i.e., how much they shifted toward a particular set of objects with a shared property. Their categorisation strategy was inferred from the sequence of objects they interacted with during free play.

First, we defined four models of infants’ categorising objects in a set with the following transitional probabilities between sequential touches: 1 = uniform play - objects touched in a random sequence, regardless of their shape or material, all object categories had a transitional probability of 1/16; 2 = shape - sets of objects touched sequentially according to their shape, ball-to-ball and cube-to-cube transitional probabilities set to 1/8, while ball-cube sequences had a probability of 0; 3 = material - sets of objects touched sequentially according to their material, soft-to-soft and hard-to-hard transitional probabilities set to 1/8, while soft-hard sequences had a probability of 0; 4 = ball bias - only balls touched sequentially to account for infants’ strong ball bias at this age, only ball-to-ball transitions had a probability of 1/4.

Next, the infants’ observed sequential interactions with the objects were compared with these transitional probability distributions using a Jensen-Shannon Divergence (JSD) dissimilarity matrix. The smaller the JSD, the closer the observed and modeled transitional probability distributions were to each other. We assessed infants’ change in JSD from pre- to post-demonstration for each of the four categorisation models. The overall index of set shifting **ShiftQ** was given by the highest differential in JSD between pre- and post-play in any categorisation model. Please note that shiftQ is a negative metric as it measures the lowest dissimilarity between modeled and observed distributions. Thus, lower shiftQ signified higher set shifting.

### Weighted phase lag index (WPLI)

The weighted Phase-Lag Index (WPLI) applied to electrophysiological signals (Canales-Johnson et al., 2021a; Olivares et al., 2025; Potash et al., 2025; Vinck et al., 2011) measures the extent to which phase angle differences between two time series *x*(*t*) and *y*(*t*) are distributed towards positive or negative parts of the imaginary axis in the complex plane. The underlying idea is that volume-conducted activity accounts for the greatest proportion of detected 0° or 180° phase differences between signals. Therefore, only phase angle distributions predominantly on the positive or negative side are considered to obtain a conservative estimate for real, non-volume conducted activity. The PLI is the absolute value of the sum of the signs of the imaginary part of the complex cross-spectral density Sxy of two real-valued signals *x*(*t*) and *y*(*t*) at time point or trial t. While the Phase-Lag Index (PLI) is already insensitive to zero-lag interactions, the WPLI further addresses potential confounds caused by volume conduction, by scaling contributions of angle differences according to their distance from the real axis, as almost ‘almost-zero-lag’ interactions are considered as noise affecting real zero-lag interactions:

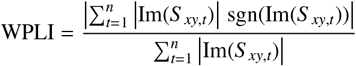

The WPLI is based only on the imaginary component of the cross-spectrum, and thus implies robustness to noise compared to coherence, as uncorrelated noise sources cause an increase in signal power. Here, WPLI was computed using the Fieldtrip toolbox (multi-taper method, fast Fourier transform, single Hanning taper, 0.5 Hz frequency resolution). Note that we used a frequency band for WPLI (i.e., 3-16 Hz) that accommodated both infant and adult spectral peaks and was comparable to the frequency ranges computed for WSMI (see below). Thus, for each epoch, we obtained a single WPLI value after computing and averaging WPLI across all electrode pairs.

### Weighted symbolic mutual information (WSMI)

We quantified the information sharing between electrodes by calculating the weighted symbolic mutual information (WSMI). This index estimates the extent to which two EEG signals exhibit non-random joint (i.e., correlated) fluctuations. Thus, WSMI has been proposed as a measure of neural information sharing (King et al., 2013) and has three main advantages. First, it is a rapid and robust estimate of signals’ entropy (i.e., statistical uncertainty in signal patterns), as it reduces the signal’s length (i.e., dimensionality) by looking for qualitative or ‘symbolic’ patterns of increase or decrease in the signal. Second, it efficiently detects highly non-linear coupling (i.e., non-proportional relationships between neural signals) between EEG signals, as it has been shown with simulated (Imperatori et al., 2019) and experimental EEG (Canales-Johnson et al., 2020a; Imperatori et al., 2019; King et al., 2013; Potash et al., 2025; Sitt et al., 2014), intracranial EEG (iEEG) (Canales-Johnson et al., 2020b), and local field potentials (LFP) (Olivares et al., 2025). Third, it rejects spurious correlations between signals that share a common source, thus prioritizing non-trivial pairs of symbols.

We calculated WSMI between each pair of electrodes, for each trial, after transforming the EEG signal into a sequence of discrete symbols defined by ordering of k time samples with a temporal separation between each pair (or τ). The symbolic transformation is determined by a fixed symbol size (κ = 3, i.e., 3 samples represent a symbol) and the variable τ between samples (temporal distance between samples), thus determining the frequency range in which WSMI is estimated (King et al., 2013). The frequency specificity *f* of WSMI is related to k and τ as follows:

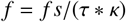

With a *fs* = 250 Hz, as per the above formula, and a *k* size of 3, this leads to a τ = 16.6 Hz. Note that we used a frequency band for WSMI that accommodated both infant and adult spectral peaks and was comparable to the frequency ranges computed for WPLI. This was done to accommodate both infant and adult theta and alpha frequencies. The weights were added to discard the conjunction of identical and opposite-sign symbols, which indicate spurious correlations due to volume conduction. The WSMI (in bits) can be calculated as:

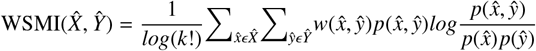

Where x and y are all symbols present in signals *X* and *Y* respectively, *w*(*x, y*) is the weight matrix, and *p*(*x, y*) is the joint probability of co-occurrence of symbol *x* in signal *X* and symbol *y* in signal *Y*. Finally, *p*(*x*) and *p*(*y*) are the probabilities of those symbols in each signal, and *K*! is the number of symbols used to normalize the mutual information (MI) by the signal’s maximal entropy. After computing and averaging WSMI across all electrode pairs, we obtained a single WSMI value for each frequency range and epoch.

### Statistical Analyses

For the ADM task analyses, we performed a Repeated Measures Analysis of Variance (RANOVA) using the three factors: Stage (Infant, Adult), Dynamics (WSMI, WPLI), and Period (Presentation, Response, Interval). In the case of the STT, RANOVA was performed using three factors: Stage (Infant, Adult), Dynamics (WSMI, WPLI), and Period (Pre, Demo, Post). Post-hoc comparisons were corrected for multiple comparisons using Tukey’s Test. Statistical analyses were performed using MATLAB (2023a) and JASP statistical software (open source).

## Acknowledgments

This research was funded by a UK Economic and Social Research Council (ESRC) Transforming Social Sciences Grant ES/N006461/1 (to V.L.), a Nanyang Technological University start-up Grant M4081585.SS0 (to V.L.), a Ministry of Education (Singapore) Tier 1 grant M4012105.SS0 (V.L.). A.C.-J. is funded by an ANID/FONDECYT Regular (1240899) and ANID/FONDECYT Regular (1251273) research grants. L.S. and S.G. are funded by the RIE2025 Human Potential Programme Prenatal/Early Childhood Grant (Award H22P0M0002) awarded to V.L., administered by A*STAR. A.C.-J. thanks Edoardo Chidichimo for his insightful comments and suggestions during the writing of this manuscript.

## Data Availability

The data and code that support the findings of this study are available from the corresponding author upon reasonable request.

## Authorship contributions

Conceptualization: L.S., S.G., A.C-J., V.L. Data analysis: L.S., S.G., A.C-J. Human recordings: L.S., S.G. Software and methods for human data: A.C-J. Visualization: A.C-J. Writing and editing: L.S., S.G., V.N., A.C-J., V.L. Funding acquisition: V.L. Supervision: V.L.

## Conflict of Interest

None declared.

## Notes

### Competing Interest Statement

The authors have declared no competing interest.

